# Proteomic profiling of advanced hepatocellular carcinoma identifies predictive signatures of response to treatments

**DOI:** 10.1101/2025.01.03.631224

**Authors:** Adèle Delamarre, Marie Decraecker, Jean-Frédéric Blanc, Sylvaine Di Tommaso, Cyril Dourthe, Jean-William Dupuy, Mélanie Moreau, Nathalie Allain, Isabelle Mahouche, Julie Giraud, Giovanni Bénard, Claude Lalou, Benoît Pinson, Paulette Bioulac-Sage, Caroline Toulouse, Audrey Morisset, Jérôme Boursier, Brigitte Le Bail, Anne-Aurélie Raymond, Frédéric Saltel

## Abstract

**Purpose:** Hepatocellular carcinoma (HCC) is the most common form of liver cancer with a bad prognosis in case of advanced HCC, only eligible for palliative systemic therapies. After a decade of exclusive sorafenib monotherapy, with a response rate of <10%, the advent of immunotherapies represents a revolution in HCC. The combination of atezolizumab/bevacizumab is recommended as the first-line systemic treatment, with a response rate around 30%. However, there are currently no predictive factors for response to these treatment options.

**Experimental Design:** We profiled, by high-resolution mass spectrometry-based proteomics combined with machine learning analysis, a selected cohort of fixed biopsies of advanced HCC. We grouped subjects according to their objective response to treatments, corresponded to a tumor regression vs tumor progression at 4 months after treatment.

**Results:** We generated a proteome database of 50 selected HCC samples. We compared the relative protein abundance between tumoral and non-tumoral liver tissues from advanced HCC patients treated. The clear distinction of these two groups for each treatment is based on deregulation for 141 protein or 87 for atezolizumab/bevacizumab and sorafenib treatment, respectively. These specific proteomic signatures were sufficient to predict the response to treatment, and revealed biological pathways involved in treatment’s resistance. Particularly, we validated a shift in tumor cell metabolism with an immunosuppressive environment involved in the resistance to atezolizumab/bevacizumab combination.

**Conclusions:** We performed an in-depth analysis of quantitative proteomic data from HCC biopsies to predict the treatment response to advanced HCC giving the ability to optimize patient management.

## Introduction

Hepatocellular carcinoma (HCC) is a major public health issue and its incidence has reached one million new cases per year worldwide (1). Alcohol consumption and an increase in obesity cases are believed to be the causes for an increased incidence of HCC of 9% per year. New cases of HCC are now primarily caused by metabolic-dysfunction-associated fatty liver disease (MAFLD) (2). HCC develops in an underlying cirrhotic liver in 80% of cases, and more rarely in chronic liver disease without cirrhosis (3).

Although screening programs diagnose approximately 40% of HCCs at a curative stage, at least 50% of patients are diagnosed at an intermediate or advanced stage (4). The prognosis remains unfavorable at these later stages due to the extensive tumor burden, the high frequency of liver dysfunction, and deteriorated health status, which limit access to curative treatments (5). Thus, HCC is the second leading cause of cancer deaths worldwide (6).

In patients with advanced HCC (Barcelona Clinic Liver Cancer [BCLC] C) or with intermediate-stage (BCLC B) disease not eligible for, or progressing despite locoregional therapies, systemic therapies are the gold standard of care (7).

Sorafenib (a multi-tyrosine kinase inhibitor, TKI) has been the standard treatment of care since 2007, based on a slightly improved overall survival (OS) in randomized controlled trials compared to supportive care (8). However, only a few patients experiment objective responses. The management of advanced HCC has recently been disrupted by the development of new effective systemic treatments, including immunotherapies that improve OS and progression-free survival (PFS). The combination of atezolizumab (anti-programmed death-ligand 1 [PDL1]) and bevacizumab (anti-vascular endothelial growth factor [VEGF]) is now the first-line standard of care for advanced HCC following the IMbrave150 study, as it increased median OS to more than 19 months in patients with preserved liver function (9–11). Moreover, a combination of an anti-PDL1 (durvalumab) and an anti-cytotoxic T-lymphocyte antigen-4 (tremelimumab) agent was approved last year after the positive Himalaya phase III trial results, strengthening the therapeutic arsenal in a first-line setting (12).

The combination of atezolizumab and bevacizumab is the most prescribed in first intent unless there is a contraindication, but there is no recommended treatment of choice among these drugs. It has been estimated that one-third of patients do not respond to the combination, without any predictive marker of (non)response (13). However, they may be potential good responders to other drugs, particularly lenvatinib (another TKI) which has a significant first-line benefit with a median OS of 19 months in a phase III trial, rather similar to survival with atezolizumab and bevacizumab (14). It has been estimated that less than 25% of patients are able to receive a second-line treatment (15). In addition, the underlying hepatic reserve and HCC progression constrain patients’ prognosis and limit access to later lines but no guidelines are available for the optimal therapeutic sequence.

In the era of sorafenib-exclusive therapy, many studies have attempted to develop tools that predict the treatment response (16–19). Some results are encouraging: a greater delay in progression has been associated with higher pre-treatment phospho-ERK levels (20). High c-Met expression predicts therapeutic efficacy (21), and non-responders express higher levels of phospho-c-Jun (22). However, none of these biomarkers have been validated in clinical practice.

Several studies have tried to predict the response to atezolizumab and bevacizumab in patients with advanced HCC using various biomarkers and signatures (13,23,24). A recent study published in *The Lancet Oncology* proposes that artificial intelligence applied on HCC digital biopsy slides could be a predictive tool for assessing sensitivity to atezolizumab/bevacizumab treatment (25). Additionally, transcriptome analysis, genomic data, and flow cytometry of peripheral blood emphasize the involvement of the immune system (particularly the immunosuppressive microenvironment) in the response to atezolizumab/bevacizumab treatment (26,27). These studies confirm the interest in identifying personalized medicine solutions for advanced HCC but still require clinical validation.

The CRAFITY (CRP and AFP in ImmunoTherapY) score has been correlated with survival and radiological response in patients receiving PD-(L)1 immunotherapy (28,29). Similarly, post-hoc analyses derived from the IMbrave150 study, have hypothesized the role of underlying liver disease in the response to immunotherapeutic treatments but the results were not confirmed by more recent clinical trials (30–33).

Thus, there is a clear and unmet need for methods to predict the treatment response and guide therapeutic selection. The proteome is the end-product of gene expression, the result of a multitude of epigenetic, transcriptional, post-transcriptional, and translational regulation, providing the functional context to cells, but no proteomic approach has been developed to predict the treatment response (34). We previously demonstrated that proteomic profiling of formalin-fixed, paraffin-embedded (FFPE) liver biopsies harbors diagnostic and prognostic value in hepatocellular adenomas (35). In this study, we compared the pre-treatment proteomic profiles of patients with advanced HCC by comparing those with an objective response to those who progressed under sorafenib and atezolizumab/bevacizumab treatment, hypothesizing that we could predict the treatment response.

## Materials and Methods

### Patient selection

We consecutively and prospectively included all patients treated with atezolizumab plus bevacizumab or sorafenib, at the University Hospital of Bordeaux and the University Hospital of Angers for advanced - unresectable and/or metastatic, not eligible for or progressing despite loco-regional therapies - HCC.

We retrospectively collected baseline characteristics of all patients before the beginning of systemic treatment, including age, sex, body mass index (BMI), performance status score (OMS), the type of underlying liver disease and cirrhosis, and the Child-Pugh and ALBI (ALbumin-BiIirubin) scores.

Cirrhosis was assumed based on clinical (signs of hepatocellular insufficiency and/or portal hypertension), biological (stigmata of hepatocellular insufficiency and/or portal hypertension), and radiological (hepatic dysmorphia, signs of portal hypertension) grounds and confirmed by liver biopsy. The diagnosis of MAFLD was made according to the 2021 international consensus (36): fatty liver highlighted by a hyperechoic aspect on ultrasound associated with obesity or overweight, type II diabetes and, in their absence, the combination of two metabolic risk factors: hypertriglyceridemia, low HDL (high-density lipoprotein) cholesterol (adjusted for sex), an increase in waist circumference (adjusted for ethnic origin), and high blood pressure.

The diagnosis of alcohol-related liver disease (ALD) was made for patients who consumed more than 21 standard doses of alcohol per week in men and 14 standard doses of alcohol per week in women without liver steatosis.

We collected data on total bilirubin, creatinine, albumin, transaminases, alkaline phosphatase, gamma-glutamyl transferase, prothrombin rate, alpha-fetoprotein (AFP), C-reactive protein (CRP), and platelets.

The tumor characteristics were the stage at the beginning of systemic treatment, the BCLC classification assessed by cross-sectional imaging, the degree of tumor differentiation, the existence of micro and/or macrovascular invasion (in cases of initial resection), the architecture of the tumor following the WHO classification, the expression levels of GS, cytokeratin 19, and the KI67 proliferation index.

All patients underwent liver biopsy with sufficient tumoral and non-tumoral tissue (> 1 mm^2^) for subsequent proteomic analysis by mass spectrometry (MS) before undergoing one of the systemic treatments.

This study was conducted while closely following the international guidance of the Good Clinical Practices and the Declaration of Helsinki. According to French national law, this study was approved by the National Ethics Committee (CPP Est III, CER-BDX 2024-106), and the Competent Authority (ANSM) was informed. The processing and use of data collected as part of this study were performed following the General Data Protection Regulation (GDPR-EU 2016/679) and complied with the Reference Methodology" (MR-004) of French Data Protection laws in force. All living participants were provided comprehensive informed notice regarding the purpose of the study, possible risks, and expected benefits, and individual participant’s rights (voluntary participation and freedom to decline participation in the trial).

Patients were included if they were 18 years or older, with histologically confirmed and radiologically measured advanced HCC.

Exclusion criteria for the study were no biopsy from a healthy liver and/or the tumor or insufficient quantity and/or quality of sample; an incomplete treatment regimen, such as less than three cycles of immunotherapy and an anti-angiogenic or more than 15 interrupted days of sorafenib use; another systemic treatment before receiving atezolizumab plus bevacizumab or sorafenib; a mixed component HCC, such as hepatocholangiocarcinoma or neuroendocrine contingent; pathological evidence obtained from a metastasis sample; an extra-hepatic recurrence.

### Follow-up

We analyzed responses and survivals of patients who received the atezolizumab plus bevacizumab or sorafenib regimen, including OS and PFS. The initial diagnostic staging was based on a computed tomography (CT) scan or magnetic resonance imaging (MRI). Treatment duration was calculated from day 1 of the regimens to the day of the disease progression, a shift to another treatment regimen, or when the patient was lost to follow-up or died. We followed the tumor response with either an abdominal CT scan or MRI, according to the Response Evaluation Criteria in Solid Tumors (RECIST; version 1.1), and the AFP level. The response was determined according to the radiological tumor assessment and/or the AFP level assessed during the first 3 months after treatment was initiated. An objective response corresponded to a > 30% tumor regression and/or a > 50% decrease in the AFP level. A > 20% increase in tumor size corresponded to an objective progression.

### Laser capture microdissection and fixation reversion

Tumoral and non-tumoral tissues (1 mm^2^) were microdissected with a PALM type 4 (Zeiss) laser micro-dissector from 5-µm-thick FFPE sections stained with hematoxylin. Each sample was replicated three times on serial sections. The proteins were extracted, the fixation reversed, and the proteins reduced and alkylated as described previously (37).

### Sample preparation for the proteomic analysis

The proteins were desalted and digested either in an SDS-PAGE gel or using the Single-pot, solid-phase-enhanced sample preparation (SP3) method (37,38).

### LC-MS/MS analysis and raw treatment mass spectrometry data

The nano-liquid chromatography-tandem mass spectrometry analysis was performed using different generations of mass spectrometers. The details of the analysis are available in **Supplementary Material**.

### Statistics and bioinformatics analysis of the proteomic data

The means of protein abundance or the ratio between responders and progressors were compared using the Wilcoxon-Mann-Whitney *U*-test. The log-rank test was used to compare survival between the biomarker-positive and biomarker-negative patients. A p-value < 0.05 was considered significant. R version 4.3.3 software (The R Foundation for Statistical Computing, Vienna, Austria) was used to perform the principal component analysis (PCA) with the R FactorMineR and factoextra packages. Missing data were imputed using the R missForest package with the R caret package.

### Integrative biology

Gene set enrichment analysis (GSEA) was performed against the Ingenuity Pathways (IPA, Qiagen, Valencia, CA, USA) database or the Gene Ontology Database Cellular Component Database. Functional pathway and interaction analyses were carried out using the IPA platform.

### Statistical analyses of the clinical data

The results are expressed as the median (interquartile range) for the quantitative data and as percentages for the qualitative data. The quantitative variables were compared using the Kruskal-Wallis test. Qualitative variables were compared using the Fisher exact test, as appropriate. A p-value < 0.05 was considered significant.

Survival analyses were estimated using the Kaplan-Meier method. The log-rank test was used to compare the survival curves established by the Kaplan-Meier method.

Univariate and multivariate survival analyses were performed using the Cox model. Variables that had p-values ≤ 0.20 in univariate analyses were integrated into the multivariate analysis.

### Three**-**dimensional cell culture

Huh7 and Hep3b cells were grown in Dulbecco’s Modified Eagle Medium (DMEM) containing 1 g/L of glucose and 10% dialyzed fetal bovine serum (FBS) and maintained in 5% CO_2_ at 37°C. The FBS was dialyzed overnight through a 3.5 kDa cut-off membrane (Thermo Fisher Scientific, #68035; Waltham, MA, USA) in a 100-fold volume of PBS to remove excess lactate. Cells were seeded in Ultra-Low Attachment 96-well plates (Corning Costar #7007; Corning, NY, USA) to form spheroids, and maintained under slight but constant agitation during 3D culture.

IACS-010759 (Abcam, #ab286961; Cambridge, MA, USA) or vehicle (DMSO 1%) was added to the culture medium 1 day after seeding. To obtain the same number of cells and the same-sized spheroids on day 3, 4,000 cells/well were seeded for the control condition vs. 8,000 cells/well for the IACS-010759-treated condition.

Peripheral blood mononuclear cells (PBMCs) from healthy donor blood were added to wells containing spheroids on day 3 (after 48 h of treatment). The PBMCs were obtained from the buffy coat by Ficoll density gradient centrifugation and then frozen. The PBMCs were thawed on the day of the experiment and counted using Trypan blue to assess viability. The PBMCs were seeded at a ratio of one tumor cell per ten PBMCs in the spheroid wells, in the same culture medium (39).

On day 5 (after 48 h of co-culture), the spheroids were washed by up and down pipetting each spheroid to remove the PBMCs that did not infiltrate. The Huh7 spheroids were collected and dissociated with 0.25% trypsin, and the cells were stained with propidium iodide (PI; Sigma, #P4864; St. Louis, MO, USA) to assess viability by flow cytometry. The Huh7 and Hep3b spheroids were collected and fixed overnight at 4°C in 4% paraformaldehyde for immunohistochemistry.

### *In vitro* metabolic analyses

The oxygen consumption rate (OCR) assessment was done for the Huh7 spheroids on day 3 (after 48 h of IACS-010759 or vehicle treatment). XFe 96-well microplates were coated with collagen type I solution according to the Campioni et al. protocol (40). Four spheroids were plated in each well in 160 μl of Seahorse XF Base Medium (DMEM medium, pH 7.4, without Phenol red and glucose, supplemented with 2 mM glutamine, 1 mM sodium pyruvate, 0.9% NaCl, and 10% FBS) and prewarmed to 37°C. The cells were incubated in a CO2-free incubator at 37°C for 1 hour. OCR was measured at baseline (basal OCR), and then successively after oligomycin (ATP synthase inhibitor), FCCP (mitochondrial uncoupling agent), and rotenone and antimycin A (inhibitors of complexes I and III, respectively) injections. OCR was normalized to protein content.

The spheroid supernatant was collected on day 3 (after 48 h of IACS-010759 or vehicle treatment) to quantify lactate. A sample of cell-free culture medium (low glucose DMEM with 10% dialyzed FBS) was used as the background control (basal lactate concentration in culture medium). The spheroid supernatants (100 µl) were extracted using an ethanol boiling method already described (41). The metabolites were separated by high-performance ionic chromatography on an Integrion chromatography station (Thermo Electron, Waltham, MA, USA) equipped with an AS11-HC 4µ column (250 × 2 mm; Thermo Electron) using a KOH gradient (42). Lactate was detected by conductometry (Dionex Integrion conductivity detector, Thermo-Electron) and quantified by comparison with pure lactate (Sigma-Aldrich #71718) standard curve. Lactate production was determined after 3 days by subtracting the lactate concentration in the spheroid supernatant from that found in the cell-free culture medium. Data for each sample were normalized to the total protein quantity in the spheroids.

### Flow cytometry

Huh7 cell spheroids were incubated for 48 h in Ultra-Low Attachment 96-well plates with CellTrace Violet-labeled PBMCs at a 1:10 ratio in complete medium (low glucose DMEM with 10% dialyzed FBS). Briefly, 1 million PBMCs were incubated with 1 μM CellTrace Violet dye (Invitrogen, #C34557; Carlsbad, CA, USA) for 20 minutes at 37°C. The cells were then incubated with complete medium for 5 minutes at room temperature and centrifuged before seeding with the spheroids. The spheroids were collected on day 5 (after 48 h of co-culture), washed, and dissociated with 0.25% trypsin. Viable cells were analyzed by side scatter and PI-negative staining. The analysis was performed on the BD FACSCanto TM II flow cytometer. Immune infiltration was assessed by quantifying the ratio of viable CellTrace Violet-positive PBMCs to total viable cells.

### Immunohistochemistry

Spheroids were collected on day 5 (after 48 hours of co-culture), washed, and fixed overnight at 4°C in 4% paraformaldehyde before the immunohistochemistry analysis. The spheroids were embedded in Histogel, dehydrated, and embedded in paraffin. Samples were cut into 3 µm sections and heated at 60°C for at least 2 hours. The paraffin was removed and the sections were rehydrated using Dako PT-link at pH 6 for 20 minutes. The slides were incubated with H_2_O_2_ for 10 minutes at room temperature to prevent endogenous peroxidase activity. Immunohistochemistry was performed using the CD45 antibody (mouse monoclonal antibody, 1:400, Dako Agilent #M070101-2; Carpentaria, CA, USA). The secondary antibody was the FLEX anti-HRP antibody and the staining procedure was performed using DAB (EnVision Flex, Dako Agilent). All images were obtained using a Nikon DS-Ri2 microscope and analyzed with QuPath v0.5.1 software.

### Statistical analyses for the *in vitro* experiments

Each value is presented as the mean ± standard error. All experiments were conducted in triplicate or more. Student’s *t*-test was employed to compare two groups. One-way analysis of variance (ANOVA) was used to compare multiple groups. A p-value < 0.05 was considered significant. Statistical analyses were performed using GraphPad Prism 10.2.2 software (GraphPad Software Inc. La Jolla, CA, USA).

## Results

### Patient selection

We first compared the tumoral proteomic profiles of 9 patients with objective responses to atezolizumab plus bevacizumab with 15 patients who progressed under the combination (**Table 1**). Most of the clinical characteristics at the beginning of the treatment were similar between the two groups, particularly median age, BMI, the disease stage (locally advanced or metastatic stage), biological test results, AFP level, macrovascular invasion, degree of differentiation, degree of proliferation, and previous loco-regional treatments. In total, 60% of the progressors had MAFLD vs. 37.5% of the responders (p = 0.10); 73.3% of the progressors had cirrhosis vs. 50% of the responders (p = 0.37); and 46.7% of progressors had an initial OMS score of 1 vs. 25% of the responders (p = 0.40). There was a more massive trabecular contingent in tumors from responders (37.5% vs. 13.3%, p = 0.38). Two patients in the responder group had ALD and hepatitis C virus (HCV) and 1 patient had ALD and hepatitis B virus (HBV).

**Table 1:** Clinical characteristics of the patients treated with atezolizumab plus bevacizumab selected for proteomic analysis. Data are numbers of patients (%) or medians (IQRs). AFP: alpha-fetoprotein; ALBI: albumin-bilirubin; ALD: alcohol-related liver disease; BCLC: the Barcelona Clinic Liver Cancer classification; BMI, body mass index; HBV: hepatitis B virus; HCV: hepatitis C virus; IQR, interquartile range. MAFLD: metabolic-associated fatty liver disease; OMS: performance status score.

| **Atezolizumab bevacizumab** | **Responders (n = 8)** | **Progressors (n = 15)** | **p-value** |
| --- | --- | --- | --- |
| **Age (years) median (IQR)** | 70.5 (61–75.5) | 73 (68.5–75.5) | 0.60 |
| **Men N (%)** | 7 (87.5) | 11 (73.3) | 0.62 |
| **Etiology N** |  |  |  |
| MAFLD | 3 (37.5%) | 9 (60%) | 0.10 |
| ALD | 4 (50%) | 2 (13.3%) |  |
| HBV | - | 1 (6.7%) |  |
| HCV | 2 (25%) | 1 (6.7%) |  |
| Healthy Liver | 1 (12.5%) | 2 (13.3%) |  |
| **Cirrhosis N (%)** | 4 (50) | 11 (73.3) | 0.37 |
| **OMS N (%)** |  |  |  |
| 0 | 6 (75) | 8 (53.3) | 0.40 |
| 1 | 2 (25) | 7 (46.7) |  |
| **Child-Pugh N (%)** |  |  |  |
| A | 7 (87.5) | 14 (93.3) | 0.42 |
| B7 | 1 (12.5) | 1 (6.7) |  |
| **ALBI score N (%)** |  |  |  |
| 1 | 3 (37.5) | 7 (46.7) | 1 |
| 2 | 3 (37.5) | 6 (40) |  |
| 3 | 1 (12.5) | 1 (6.7) |  |
| **AFP (ng/mL) median (IQR)** | 8.9 (4.7–6176) | 14.45 (7–4824.50) | 0.71 |
| **BCLC stage N (%)** |  |  |  |
| BCLC B | 3 (37.5) | 5 (33.3) | 1 |
| BCLC C | 5 (62.5) | 10 (66.7) |  |
| **Degree of differentiation (%)** |  |  |  |
| Well-differentiated | 4 (50) | 4 (26.7) | 0.39 |
| Moderately/poorly differentiated | 4 (50) | 10 (66.7) |  |
| **Vascular microinvasion (%)** | 2 (25) | 5 (33.3) | 0.36 |
| **Macro trabecular and massive contingent (%)** | 3 (37.5) | 2 (13.3) | 0.38 |
| **Previous treatment (%)** | 4 (50) | 8 (53.3) |  |
| Surgery | 2 | 4 | 0.69 |
| Chemoembolization | 3 | 4 |  |
| Radioembolization | - | 2 |  |
| Radiofrequency | - | 2 |  |

We compared the tumoral proteomic profiles of 9 patients with objective responses to sorafenib with 7 patients who progressed under sorafenib (**Table 2**). Most of the characteristics at the beginning of the treatment were similar between the two groups. More ALD patients were observed in the responder group (66.7% vs. none in the progressor group, p = 0.42), and they more frequently had cirrhosis (77.8%% vs. 33.9%, p = 0.3). Progressors had a worse performance status score (OMS = 1 for 28.6% vs. none in the responder group). Two patients who did not respond to sorafenib had a massive trabecular contingent (28.6% vs. none of the responders; p = 0.41).

**Table 2.**
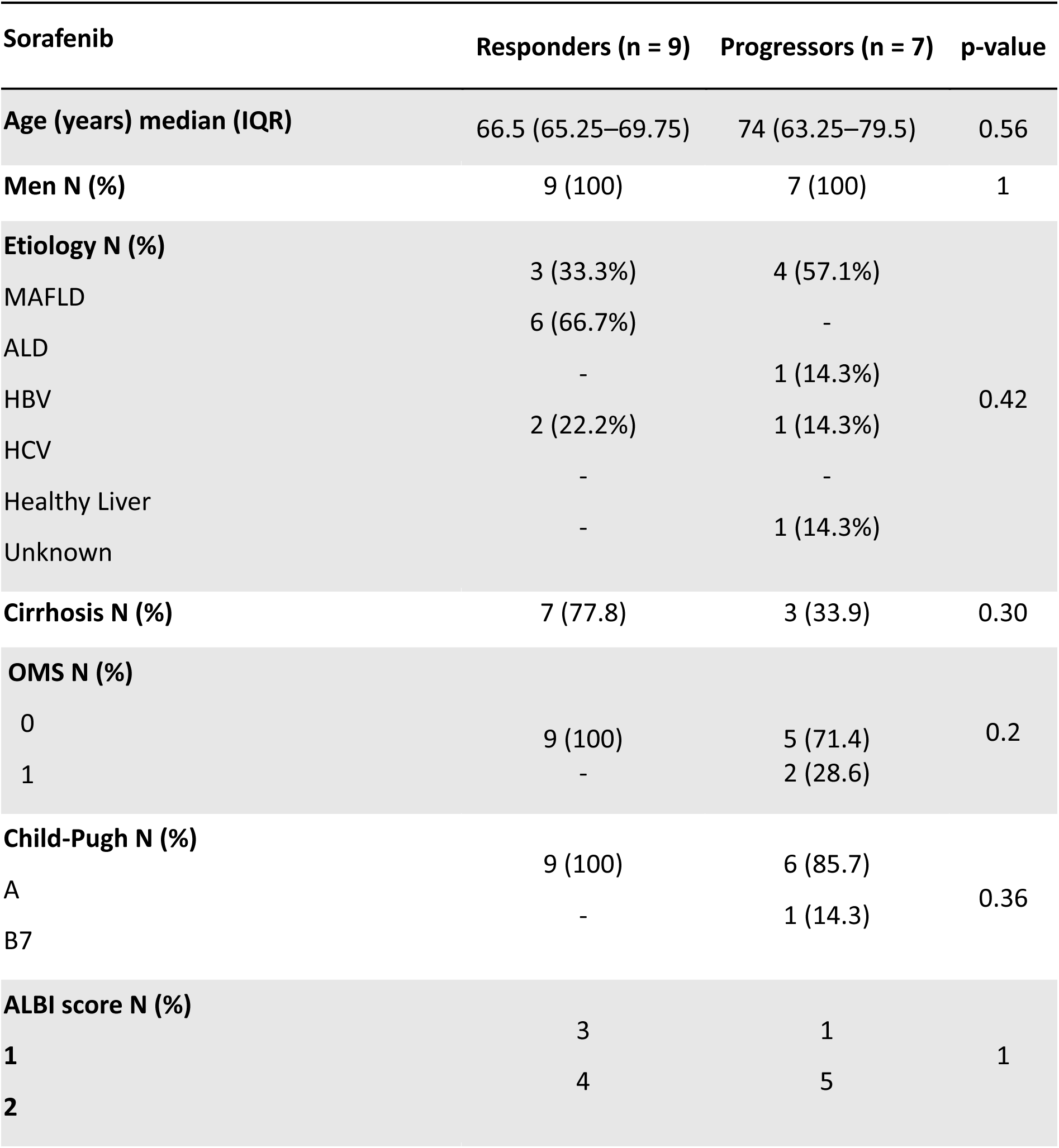
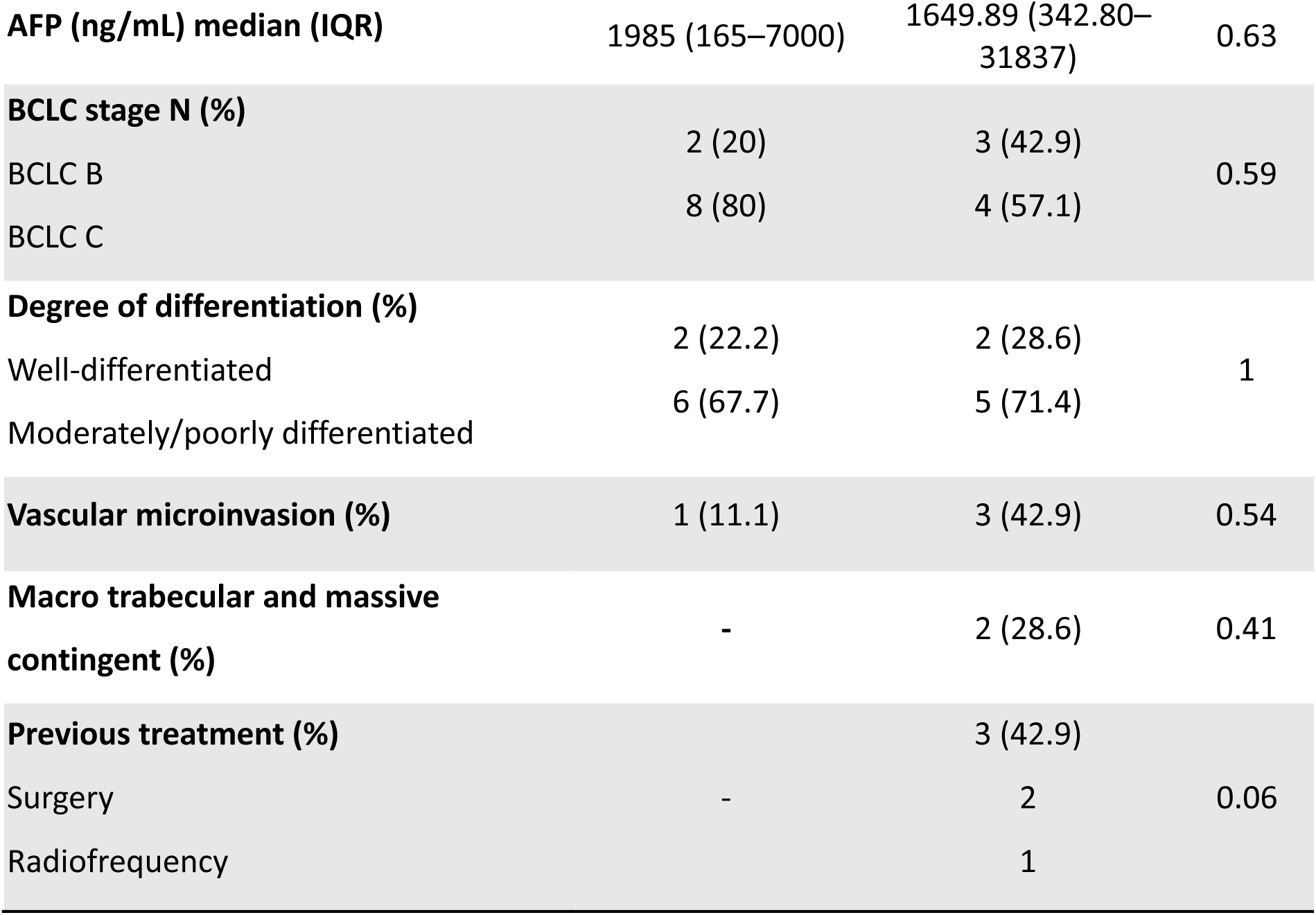
Clinical characteristics of the patients treated with sorafenib and selected for proteomic analysis. Data are numbers of patients (%) or medians (IQRs). AFP: alpha- fetoprotein; ALBI: albumin-bilirubin; ALD: alcohol-related liver disease; BCLC: the Barcelona Clinic Liver Cancer classification; BMI, body mass index; HBV: hepatitis B virus; HCV: hepatitis C virus; IQR, interquartile range. MAFLD: metabolic-associated fatty liver disease; OMS: performance status score.

Two patients in the responder group had ALD and HCV. Three patients who did not respond to sorafenib had been previously treated by surgery (N = 2) or radiofrequency (N = 1) for a localized HCC.

### Clinical outcomes

The median OS was significantly longer in responders to atezolizumab plus bevacizumab than in progressors (median OS, 28.37 (24.8–NA) months vs. 6.83 (5.4– 15.1) months respectively, p = 0.00037; **Suppl. Figure 1**). The median OS was numerically longer in responders to sorafenib than in progressors (median OS, 12.2 (9.97–NA) months vs. 4 (3.07–NA) months, respectively, p = 0.21; **Suppl. Figure 2**). PFS was significantly longer in responders to atezolizumab plus bevacizumab than in progressors (median PFS, 6.75 (6.0–NA) months vs. 1.5 (1.5–3) months respectively, p < 0.0001; **Suppl. Figure 1**). Similarly, PFS was significantly longer in responders to sorafenib than in progressors (median PFS not reached (6.0–NA) vs. 1.5 (NA–NA) months respectively, p = 0.00018; **Suppl. Figure 2**).

### Biopsy proteomic profiling differentiates patients with an objective response from those with tumor progression using atezolizumab/bevacizumab

After selection, FFPE biopsies were used to extract the patient’s proteome. First, pathologists defined the areas of interest for laser microdissection to ensure a uniform composition of liver and tumor tissue, avoiding non-relevant areas, such as necrotic tissue or vessels. The same area corresponding to 1 mm^2^ was captured from the tumor and adjacent non-tumor tissues. Then, we compared the relative protein abundance levels between the tumor (T) and adjacent non-tumoral tissue (NT) from diagnostic biopsies before treatment (**Figure 1a**). The set of proteins whose protein expression was dysregulated between T and NT corresponded to the individual tumor proteomic profile for each patient as defined by Dourthe et al. (35). We identified and quantified an average of 3,227 proteins/patient (with > 2 specific peptides). We then compared the tumor proteomic profiles of patients who responded with those who had progressed under treatment. A total of 141 proteins were significantly differentially deregulated between responders and progressors under the atezolizumab/bevacizumab treatment. Based on these 141 proteins, we tested the possibility of discriminating between patients according to their responses. PCA separated the two groups of objective response vs. progression (**Figure 1b**). As shown in the heatmap of this dataset, the intragroup proteomic profiles were heterogeneous but this 141-based protein signature distinction between the two responding groups (**Figure 1c**).

**Figure 1:**
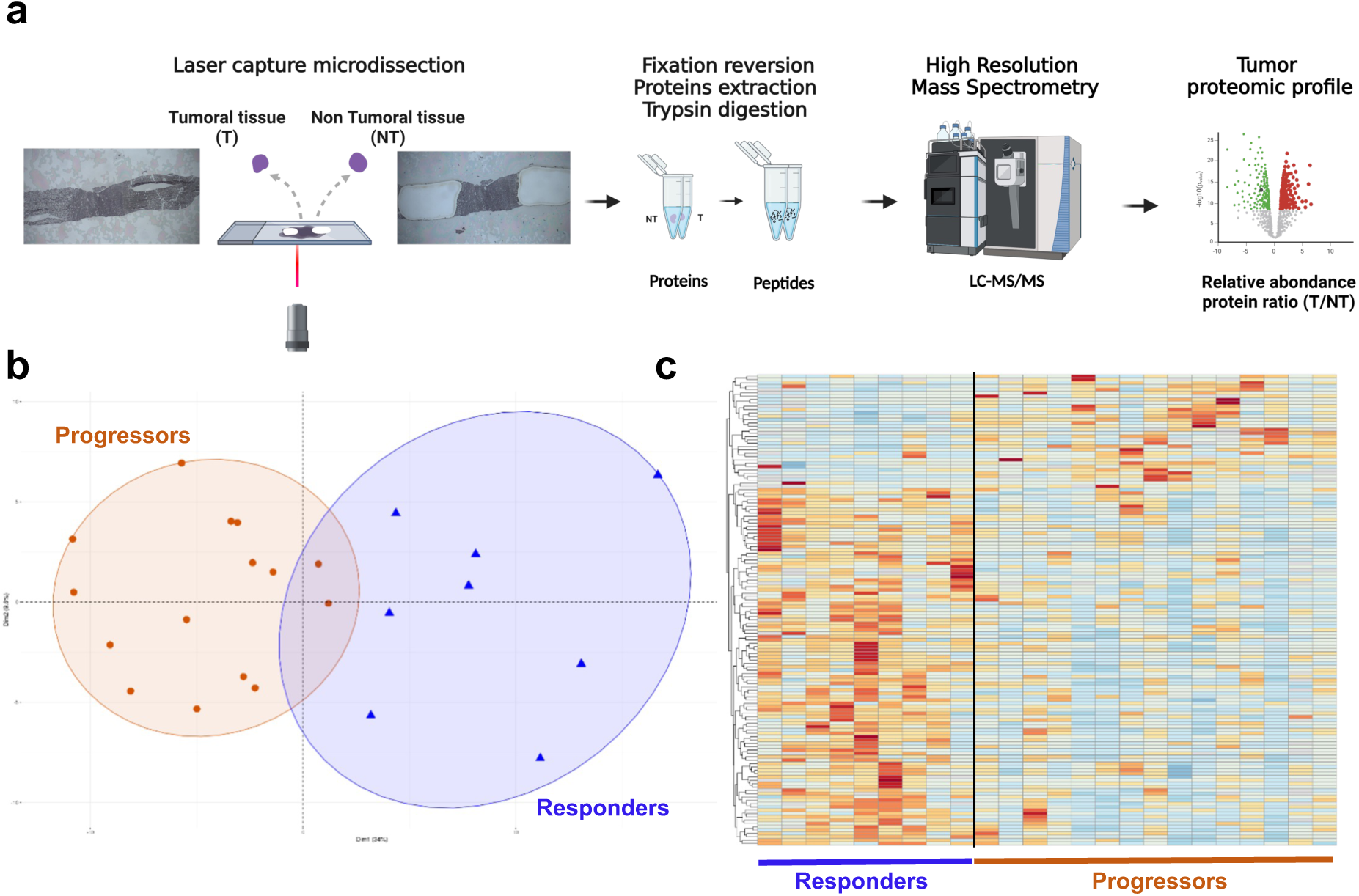
Proteomic profile of biopsies before treatment to discriminate between responders and progressors using the atezolizumab/bevacizumab combination. (a) Workflow analysis for the proteomic profiling of tissue. (b) Principal component analysis of the proteomic profiles showing deregulated protein expression between tumoral vs non-tumoral (T/NT) tissues in patients with an objective response (responders n = 9) or with progression (progressors n = 15) using atezolizumab/bevacizumab. The dataset was reduced to a signature of 141 proteins with significantly different expression between patients as responders (orange) or progressors (blue). (c) Heatmap of the 141 proteins that were significantly differentially expressed between responders and progressors for each patient.

### Analysis of the biological pathways involved in the atezolizumab/bevacizumab response

The majority (72%) of discriminating proteins were related to mitochondria according to the GSEA against the Gene Ontology (GO) Cellular Component database (**Figure 2a, 2b**). A significant decrease in the T/NT ratios of the proteins belonging to mitochondrial respiration was observed in the progressor group: oxidative phosphorylation where all mitochondrial respiratory chain complexes are affected as a whole (**Figure 2c and d**), and the beta-oxidation pathway of fatty acids (**Figure 2c**). The same trend was found for TCA cycle proteins but without statistical significance (**Figure 2c**). When we analyzed more precisely the different mitochondrial complexes, we highlighted that all were impacted in progressor’s patients (**Figure 2d**).

**Figure 2:**
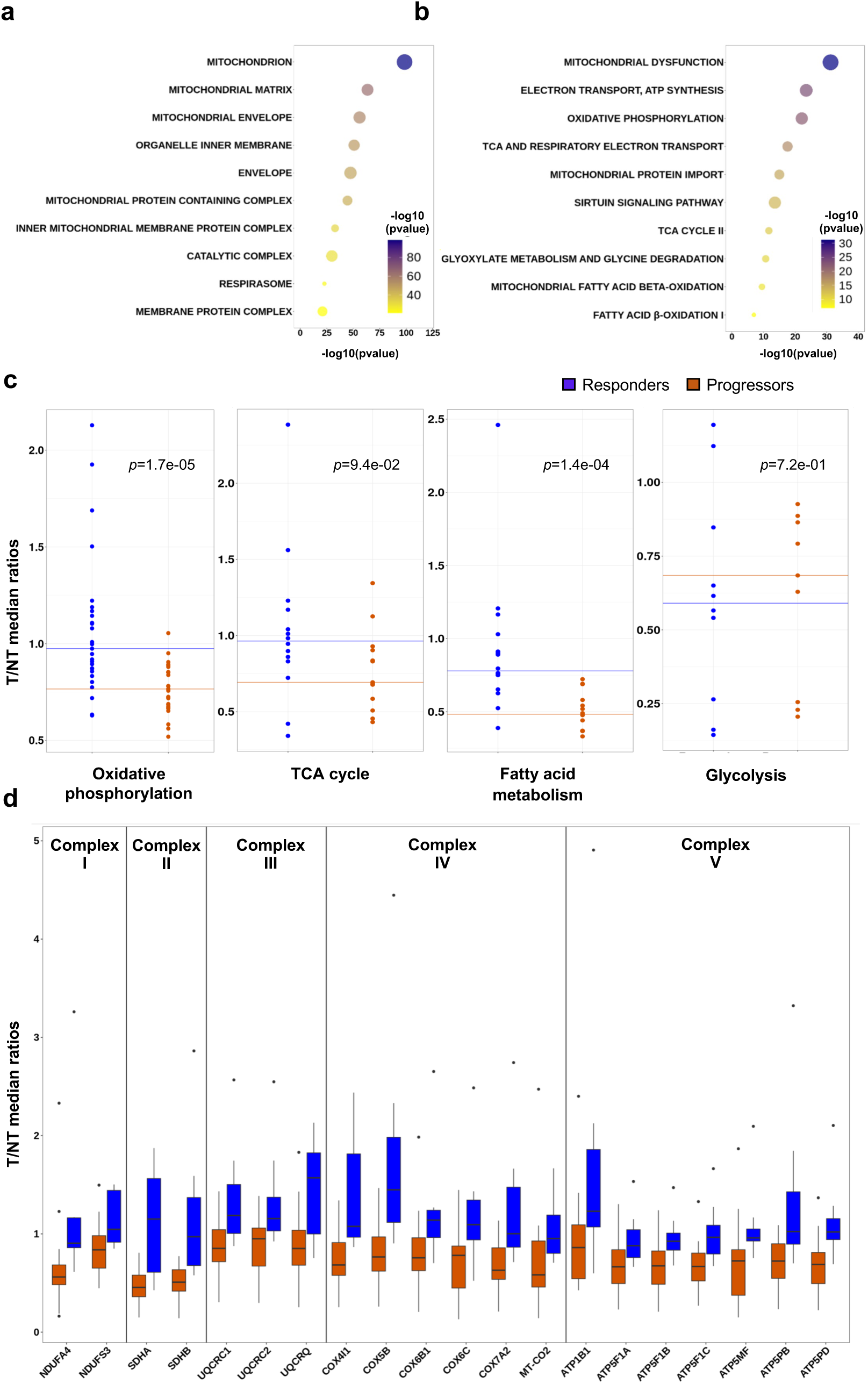
Biological pathways associated with energy metabolism. **(a-b)** Gene Set Enrichment Analysis obtained from the Gene Ontology (GO) Cellular Components (**a**) and the Ingenuity Pathways (IPA, Qiagen, canonical pathways) database (**b**) using the 141 proteins that were significantly differentially expressed between patients with an objective response versus progression using atezolizumab/bevacizumab. **(c)** Median tumoral (T)/non-tumoral (NT) ratios for proteins associated with energy metabolic pathways (oxidative phosphorylation, Krebs cycle, fatty acid oxidation, and glycolysis) for each atezolizumab/bevacizumab response group. **(d)** Details of the T/NT ratios OF oxidative phosphorylation enzymes for each mitochondrial respiratory chain complex (pathways created with Ingenuity Pathways (IPA Qiagen).

The numerous expression ratios of the glycolytic proteins tended to be higher in the progressors’ biopsies without reaching significance (**Figure 2c**).

### Inhibiting oxidative phosphorylation modulates immune cell tumor infiltration in 3D HCC models

Exploratory proteomic data have shown a downregulation of Oxphos proteins in the mitochondrial respiratory chain, which, under physiological conditions, produce energy through substrates provided by the TCA cycle and the fatty acid beta-oxidation pathway. A metabolic shift in tumor cells called the Warburg effect occurs at the expense of the Oxphos pathway and quickly provides high energy for cancer proliferation, converting pyruvate to lactate through aerobic glycolysis, even in the presence of oxygen (43). The lactate is excreted out of the cell by monocarboxylate transporters, contributing to extracellular acidification and immune suppression (44). We hypothesized that the metabolic reprogramming identified using proteomic profiling, which is a hallmark of cancer, could be a mechanism of resistance to atezolizumab and bevacizumab in HCC patients (45).

To validate this hypothesis, we developed a 3D model to mimic the metabolic shift in tumor cells that decreased oxidative phosphorylation, and studied immune cell infiltration into the tumor (**Figure 3a**). This metabolic shift identified *in vivo* was mimicked in a spheroid model of HCC cell lines (Huh7 and Hep3b) using a specific and highly potent inhibitor of mitochondrial respiratory chain complex I called IACS-010759 (46). As all respiratory chain complexes in the proteomic data were affected, we chose to inhibit complex I, which is the main entry point for electrons into the mitochondrial respiratory chain and plays a major role in mitochondrial respiration. IACS-010759 activity was assessed using Seahorse’s measurement of the oxygen consumption rate (OCR) (**Figure 3b** **and** **S3a**). A 5 nM dose resulted in a 50% decrease in basal OCR and was selected for the following experiments.

**Figure 3:**
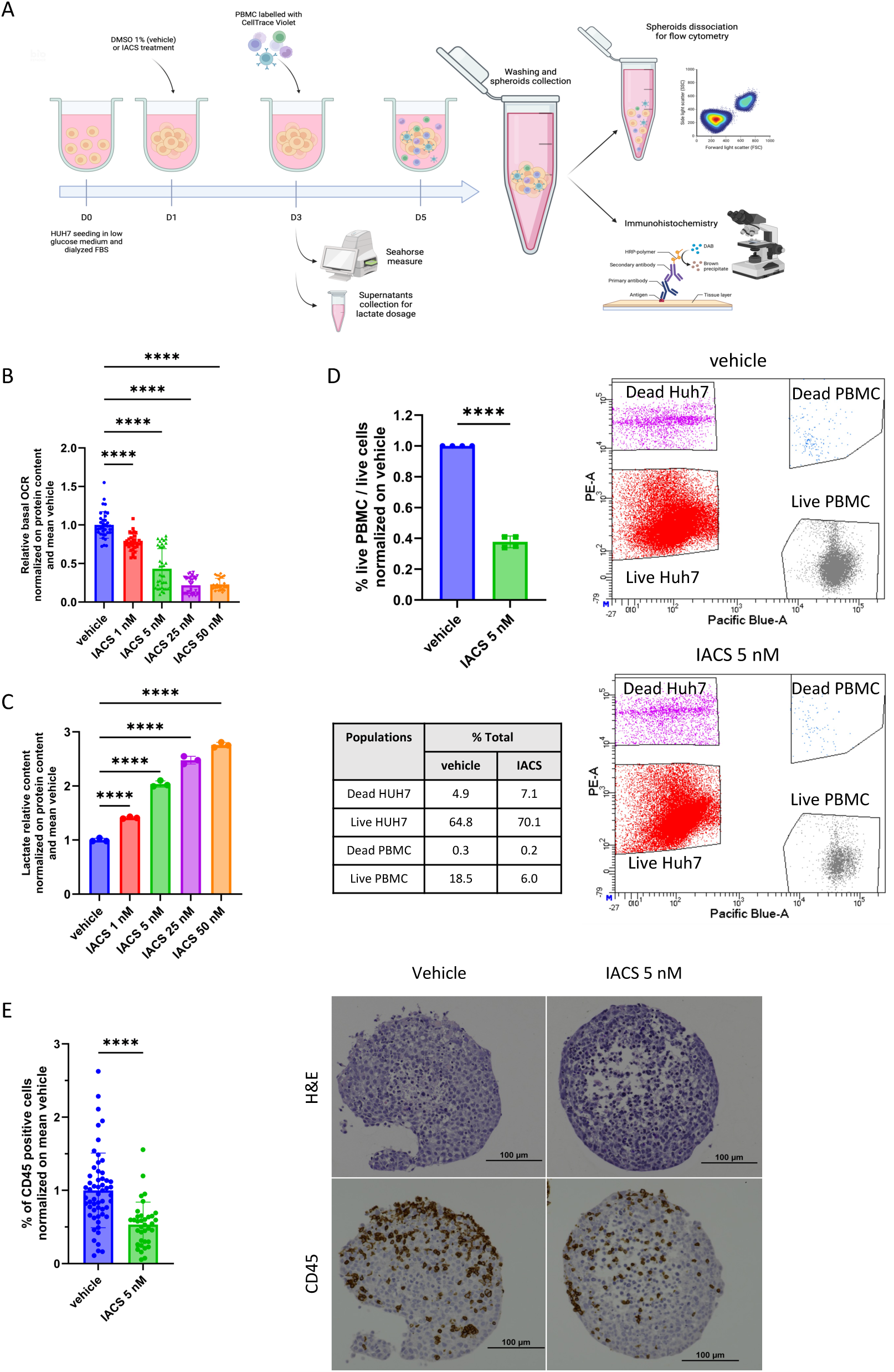
Tumor energy metabolism status modulates the infiltration of immune cells. **(a)** Experimental plan created with BioRender.com. **(b)** Relative basal OCR of the Huh7 spheroids after 48 h of IACS-010759 treatment, normalized to the protein content and mean vehicle (mean basal OCR in vehicle: 219.58 pmol/min). **(c)** Relative lactate production in Huh7 spheroid supernatants after 48 h of the IACS-010759 treatment, normalized to protein content and mean vehicle (mean lactate concentration in the vehicle: 1.60 mM). **(d)** Immune infiltration of PBMCs in Huh7 spheroids was assessed by flow cytometry normalized to the mean vehicle (mean percentage of live PBMCs to total live cells in the vehicle: 32.13%), with representative flow cytometry data. **(e)** Immune infiltration of PBMCs into Huh7 spheroids assessed by CD45 staining normalized to the mean vehicle (mean percentage of CD45 positive cells in the vehicle: 26.04%), with representative H&E and CD45-stained photographs. Data are mean ± SEM. n = 3, unpaired *t*-test or one-way ANOVA for multiple comparisons, **** p < 0.0001, *** p < 0.001, ** p < 0.01, * p < 0.05.

We assessed lactate production as a read-out of the metabolic shift from oxidative metabolism to glycolysis. Lactate production was measured in Huh7 spheroid supernatants by high-performance ion chromatography and conductimetry. The IACS- 010759 treatment resulted in a dose-dependent increase in lactate production with a two-fold increase at 5 nM (**Figure 3c**).

We assessed the effect of HCC spheroid treatment with IACS-010759 on immune infiltration through the co-culture of Huh7 spheroids and PBMCs. Immune infiltration was quantified by flow cytometry using the ratio of live PBMCs/all live cells. The IACS- 010759 treatment resulted in a 60% reduction in immune infiltration into the spheroids (**Figure 3d**). A control experiment was performed to assess the effect of IACS-010759 on PBMC infiltration capacity and showed that pre-treating PBMCs with IACS-010759 did not affect the infiltration of PBMCs into the Huh7 spheroids (**Figure S3b**).

We confirmed these results by immunohistochemistry using the CD45 immune marker to visualize and quantify immune cell infiltration. Immune infiltration was quantified using the percentage of CD45-positive cells. The IACS-010759 treatment resulted in a 50% reduction in immune infiltration in the Huh7 spheroids (**Figure 3e**) and a 25% reduction in the Hep3b spheroids (**Figure S3c**).

### Specificity of the tumoral proteomic profile to predict the treatment response

Then, we assessed the specificity of energy metabolic protein deregulation of atezolizumab/bevacizumab resistance in patients treated with another treatment. Using the same strategy, we compared the proteomic profiles of responders with progressors under the sorafenib treatment. We identified and quantified 3,153 proteins/patient (with > 2 specific peptides per protein), allowing us to extract 87 proteins differentially, and significantly deregulated between responders and progressors. PCA differentiated the patients according to their response to the TKI (**Figure 4a**). The heatmap of this dataset shows the distinct proteomic profiles between patients with an objective response and the progressors (**Figure 4b**). Only four common proteins were observed between the signatures, demonstrating the specificity of the proteomic profiles for each treatment (**Figure 4c**).

**Figure 4:**
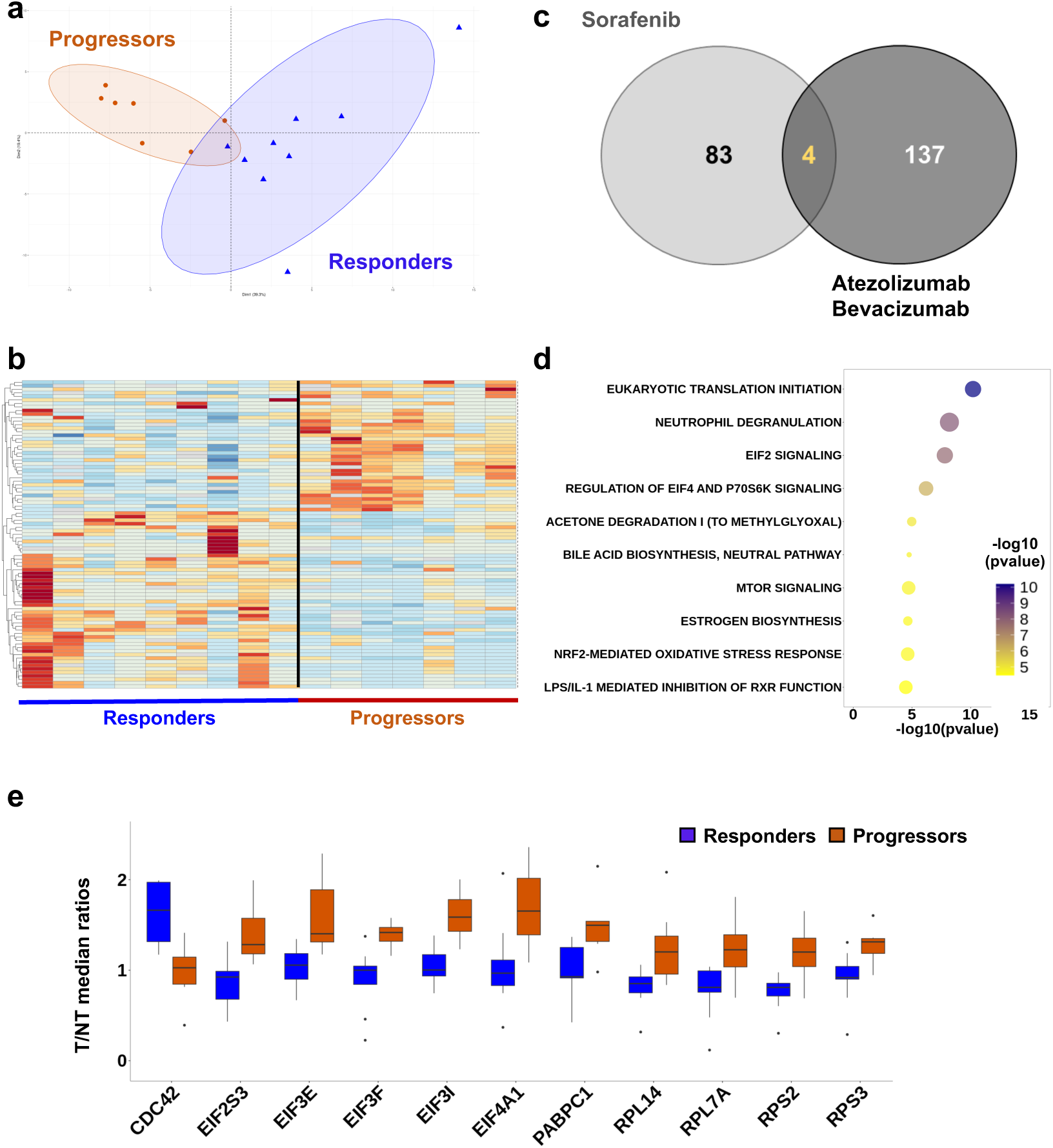
Proteomic profile of biopsies before treatment to discriminate between responders and progressors using sorafenib. **(a)** Principal component analysis of the proteomic profiles showing deregulated protein expression between tumoral vs. non-tumoral (T/NT) tissues in patients with an objective response (n = 9) or with progression (n = 15) under the sorafenib treatment. The dataset was reduced to a signature of 87 proteins with significantly different expression levels between progressors (orange) and responders (blue). **(b)** Heatmap of the 87 proteins that were significantly differentially expressed between the two groups**. (c)** Comparison of proteins whose expression discriminated the responses (responder or progressor) to the two treatments (atezolizumab/bevacizumab or sorafenib). **(d)** Gene Set Enrichment Analysis was obtained from the Ingenuity Pathways (IPA, Qiagen, canonical pathways) database using the 87 proteins that were significantly differentially expressed between patients with an objective response vs. those who progressed using sorafenib. **(e)** Details of T/NT ratios of proteins involved in protein translation and the mTOR pathway.

We performed biological pathways enrichment analysis by GSEA and did not detect any enrichment of energy metabolic pathways. The protein translation pathways (translation, eukaryotic initiation factor [eIF] signaling) and mTOR signaling were significantly upregulated in sorafenib progressors, while cytochrome activities were significantly downregulated, concordant with some published data, validating our approach (**Figure 4d and 4e**) (47–53). Interestingly, we identified an original target named Cdc42, a Rho-GTPase involved in cancer cell invasion, which was here significantly overexpressed in responders (T/NT median ratio = 1.95) and not modified in progressors (T/NT median ratio = 1.02) (**Figure 4e**) (54). These targets also include proteins involved in translation, which were systematically overexpressed by an average of 40%. In particular, we noticed three proteins of the EIF3 complex (EIF3E, F and I), a complex involved in tumorigenesis (**Figure 4e**) (50).

These data demonstrate the potential and the specificity of proteomic profiling to predict the response to treatments, and also for identifying pathways involved in treatment resistance.

## Discussion

The lack of markers to predict the response to treatments in advanced HCC remains a big challenge, so we must strive to determine the optimal treatment and therapeutic sequence, particularly for newly available combinations. We demonstrated for the first time that the information required for this predictive strategy was available in the proteome of an advanced HCC diagnostic stage biopsy, using laser capture and high-resolution mass spectrometry of small FFPE liver samples. We found distinct proteomic profiles associated with the efficacy of atezolizumab plus bevacizumab and sorafenib, compared to non-responders. Interestingly, responders to this combination exhibited a different tumoral proteomic profile to that of responders to sorafenib, suggesting the potential specific theragnostic value of proteomic profiling.

Our biological validation further highlights the distinct mechanisms involved in treatment resistance, and our functional analyses showed that a reconditioned energy metabolism with a lower oxidative phosphorylation rate played an important role in the resistance to atezolizumab plus bevacizumab.

The expression of glycolysis enzymes was slightly increased in patients who progressed under treatment. Only hexokinase 1 was significantly overexpressed between the two response groups. This discrete increase in the expression of glycolytic enzymes is probably associated with an increase in glycolytic flux, and should be able to compensate for the drop in mitochondrial respiration to supply tumor cells.

The metabolic reprogramming of tumor cells, known as the Warburg effect, is a major hallmark of cancer and is well described for other types of cancers (55–59). We have demonstrated *in vitro* that by inhibiting the Oxphos pathway, the decrease in mitochondrial respiration could lead to a significant decrease in immune cell infiltration into spheroid tumors, which is a primary step in immune escape, another fundamental hallmark of cancer (60). This mechanism, through the production of lactate and environment acidification, contributes to an immunosuppressed environment and leads to the escape of tumors from immunosurveillance.

We identified biological pathways in the proteomic profile known to be involved in resistance to sorafenib, and which served as a positive control for our approach. The PI3K/AKT/mTOR signaling pathway regulates crucial cellular processes in physiological settings, such as proliferation, survival, metabolism, motility, and angiogenesis, and HCC is frequently associated with changes in mTOR signaling (47). HCCs with upregulated PI3K/AKT signaling are more aggressive and at risk for earlier recurrence. eIF pathways are involved in initiating protein translation and eiF3 (a 13- subunit complex), in particular, interacts with the PI3K/AKT/mTOR pathway through phosphatidylinositol-3-kinases to initiate translation (48,49). High expression of certain eIF3 subunits has been associated with proliferation, invasion, and tumorigenicity in HCC (50,51). In this study, increased expression of translational actors in patients’ progressors could be the consequence of a reprogramming tumor proteome. Characterizing this translational reprogramming could reveal new pharmacological targets to counter sorafenib resistance.

Further, it will be important to ascertain whether tumor energy status similarly affects the response to durvalumab/tremelimumab, an immunotherapeutic combination recently used for advanced HCC (12). The absence of anti-angiogenic VEGF induces specific features of the predictive factors of the response. Moreover, the different proteomic profiles between responders to atezolizumab plus bevacizumab and sorafenib offer additional insight into selecting treatments. Further experiments will be fundamental to assess the distinct mechanisms involved in each class of treatment responses.

Challenges remain, including ensuring treatment adherence and optimizing dosages, particularly with TKIs, such as sorafenib, known for their adverse effects. While the proteomic profiles were notably consistent among the responders, there was less distinctiveness among non-responders, possibly due to challenges in ensuring adherence to sorafenib. Sorafenib is not very tolerable. It has been linked to several adverse events and often leads to treatment interruptions or dose adjustments. Interpreting the signatures must consider treatment observance, and we prospectively confirmed optimal dosing with fewer than 15 days of interruption; this relied on subjective patient reporting in this real-world study.

Efforts are now focused on identifying non-invasive response predictive markers using artificial intelligence and other means such as blood biomarkers, circulating DNA, circulating cells, and exosomes. As non-invasive strategies are not yet available, and identifying a unique pathway or enzyme does not fully elucidate the resistance to treatment in patients with HCC, we have easy access to a vast amount of information in liver biopsies, highlighting the importance of larger proteomic signatures for a theragnostic purpose. The proteome is the end-product of a multitude of regulated gene expression, including transcriptional, translation, and post-translational processes. It reflects cellular functionalities and consequently is the most representative of the biological functions involved in the tumor phenotype. Analyzing the tumor’s and the adjacent liver’s proteomes allowed us to access valuable information about the tumor microenvironment.

Our unique method allowed us to define the tumor proteomic profile of each patient in the form of a tumor identity card. In the future, this tool can be used to choose the most suitable treatment for each patient by comparing it to a database of reference profiles, choosing the treatment for which the tumor profile has the most similarities with those of patients who had an objective response.

There is an unequivocal need to establish theragnostic signatures for HCC treatments in larger cohorts to enhance their validity. However, even in a small subgroup of patients, our methodology demonstrated that we could correctly differentiate patients according to their response and that we could reveal biological pathways involved in treatment resistance. Proteomic profiling has the potential to improve patients’ outcomes. The insight gained from this study will inform future efforts in developing clinical applications for prediction and, thus, personalized management of patients, opening the way to precision medicine for advanced HCC.

## Grant support

This work was supported by the Nouvelle Aquitaine Region (European FEDER Funds), the Aquitaine Science Transfert, the SIRIC BRIO, and the Arc Foundation. Sylvaine Di Tommaso, Cyril Dourthe, and Anne-Aurélie Raymond were supported by the Nouvelle Aquitaine Region (European FEDER Funds) and the HepA association. Cyril Dourthe was supported by cancéropôle grand sud-ouest. Sylvaine Di Tommaso and Cyril Dourthe were supported by Arc Foundation (Arc Sign’it program). The Integrion chromatography station used for lactate quantification was purchased with the financial support of both the SIRIC BRIO (COMMUCAN) and the Region Nouvelle-Aquitaine (AAPPF2021-2020-12000110).

The data that support the findings of this study are available from the corresponding author upon reasonable request.

We did not reproduce any material (fragments of text, tables, figures) from other sources.

We hereby certify that Textcheck has checked and corrected the English in the manuscript named above (REF 24060405).

## Potential conflicts of interest

JFB: Bayer, ESAI, IPSEN, ROCHE, ASTRA-ZENECA, BMS

MD: Roche, Servier

## Author Contributions

Study concept and design: AD, MD, GB, AAR, FS, JFB Manuscript writing: AD, MD, AAR, FS, JFB

Funding was obtained by: AAR, FS, JFB

Patient sample collection: MD, CT, JFB, BLB, AM, JB

## Data acquisition

AD, MD, MM, NA, CL, BP (*in vitro* analyses), BLB, PBS (histopathology interpretation), SDT (laser microdissection and sample preparation); JWD (mass spectrometry analysis and raw mass spectrometry data processing), CD (proteomics data post-processing, statistical analysis and data formatting); AAR (biological interpretation of proteomics data).

All the authors reviewed the final version and approved for submission.

## Acknowledgements

The authors thank Drs Mireille Simon, MD (Hospital of Pau) and Franck Audemar, MD (Hospital of Bayonne) for the help in patients’ recruitment.

The authors thank S. Vazquez from the Metabolic Analyses Service from the TBMCore Service Unit (CNRS UAR3427 INSERM US05 University of Bordeaux) for lactate quantification, the members of the FACSility plaform (TBMcore) for flow cytometry experiments and the members of the Histopathology platform (TBMCore) for immunohistochemistry experiments.

## Abbreviations

AFP: alpha-fetoprotein;
ALBI: albumin-bilirubin;
ALD: alcohol-related liver disease;
BCLC: Barcelona Clinic Liver Cancer classification;
BMI,: body mass index;
CK-19: cytokeratin 19;
CRP: C-reactive protein;
CTV: CellTrace Violet;
DMEM: Dulbecco’s Modified Eagle Medium;
EIF: eukaryotic initiation factor;
FFPET: formalin-fixed and paraffin-embedded tissue;
GGT: gamma-glutamyl transferase;
GO: Gene Ontology;
GS: glutamine synthetase (GS);
HBV: hepatitis B virus;
HCC: hepatocellular carcinoma;
HCV: hepatitis C virus;
HDL: high-density lipoprotein;
IQR,: interquartile range;
KI-67: index of proliferation;
LC-MS/MS: liquid chromatography-tandem mass spectrometry;
NT: non-tumoral;
MCT: monocarboxylate transporters;
MS: mass spectrometry;
MAFLD: metabolic dysfunction-associated fatty liver disease;
MRC: mitochondrial respiratory chain;
OCR: oxygen consumption rate;
OMS: performance status score;
ORR: objective response rate;
OS: overall survival;
PAL: alkaline phosphatase;
PBMCs: peripheral blood mononuclear cells;
PDL1: programmed death-ligand 1;
PI: propidium iodide;
PFS: progression-free survival;
PT: prothrombin rate;
SEM: standard error;
T: tumoral;
TKI: tyrosine kinase inhibitor;
VEGF: vascular endothelial growth factor;
WHO: World Health Organization.

---

**Supplementary figure 1.**
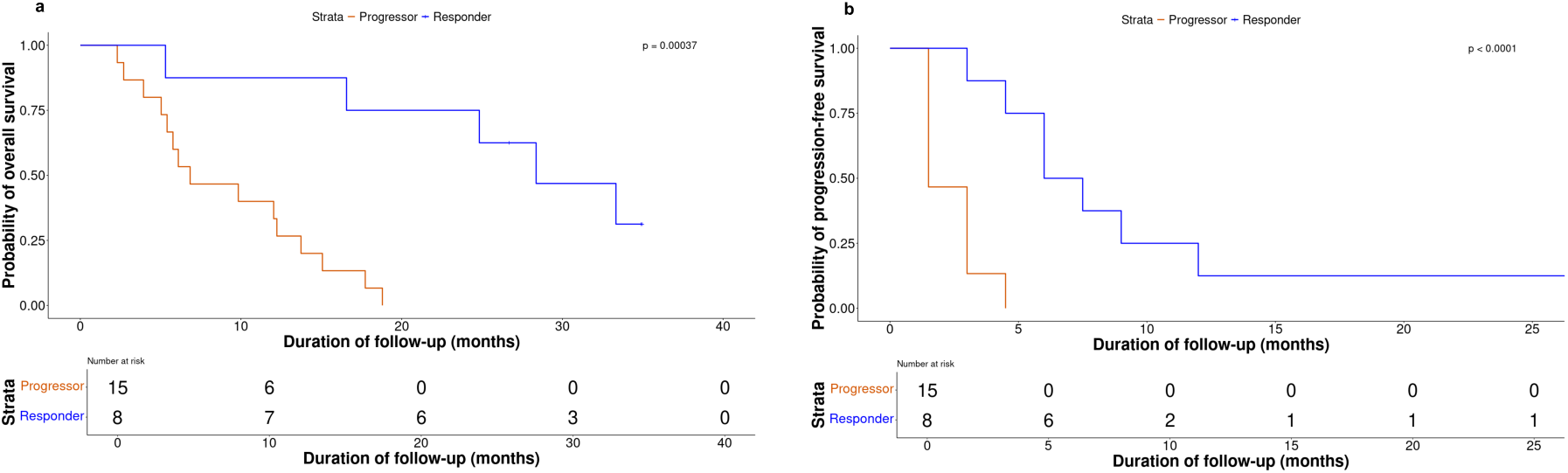
Outcomes in the atezolizumab plus bevacizumab population according to their treatment responses. **(a)** Overall survival; **(b)** Progression-free survival. Patients who progressed under the combination are shown in red; patients who responded to the combination are shown in blue. Survival curves were created using the Kaplan-Meier method, and the log-rank test was used to compare the curves. A p-value < 0.05 was considered significant.

**Supplementary figure 2.**
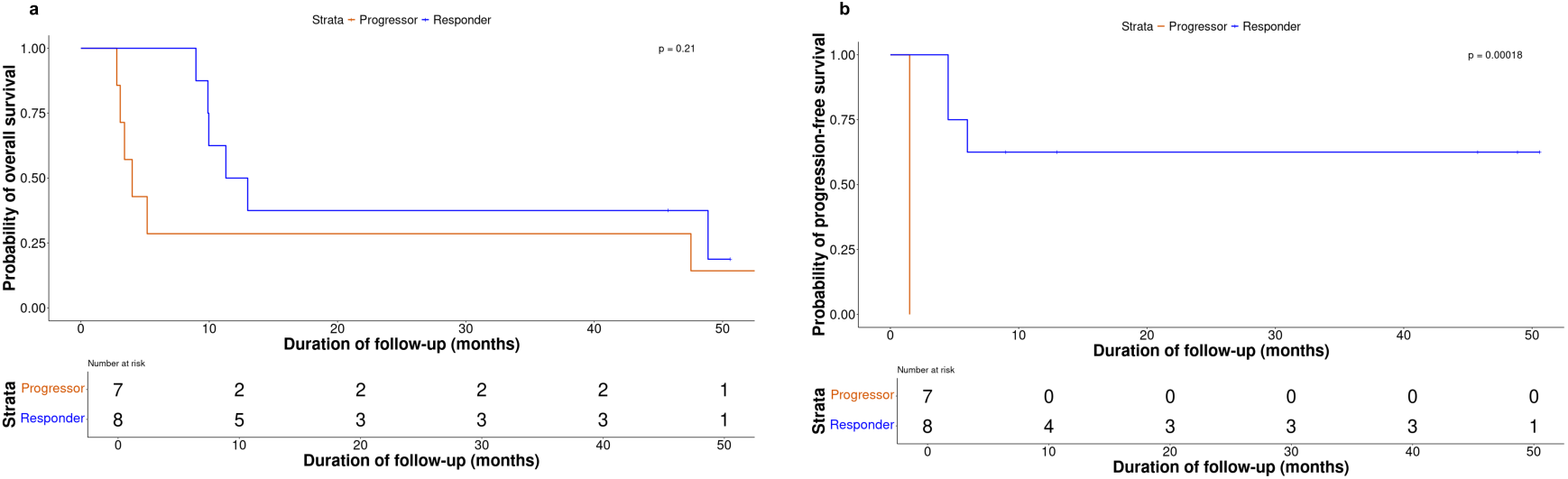
Outcomes in the sorafenib population according to their treatment responses. **(a)** Overall survival; **(b)** Progression-free survival. Patients who progressed under sorafenib are shown in red; patients who responded to sorafenib are shown in blue. Survival curves were created using the Kaplan-Meier method, and the log-rank test was used to compare the curves. A p-value < 0.05 was considered significant.

**Supplementary figure 3.**
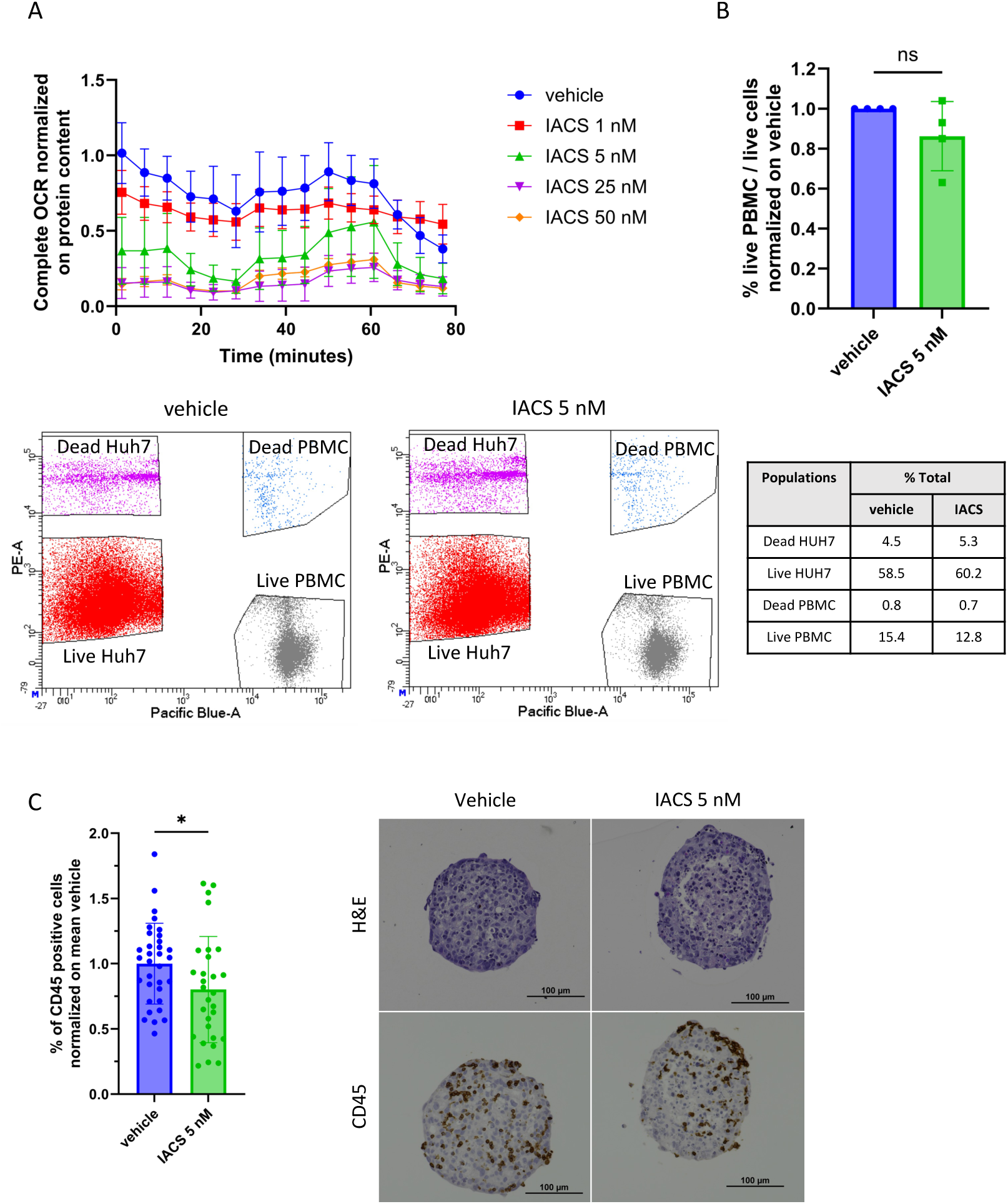
Tumor energy metabolism status modulates the infiltration of immune cells. **(a)** Representative Seahorse measure of the complete OCR of the Huh7 spheroids after 48 h of IACS-010759 treatment, normalized to protein content. **(b)** Immune infiltration of PBMCs pre-treated with IACS-010759 for 48 h in Huh7 control spheroids assessed by flow cytometry, normalized to the mean vehicle (mean percentage of live PBMCs of the total live cells in the vehicle: 37.52%), with representative flow cytometry data. **(c)** Immune infiltration of PBMCs in the Hep3b spheroids assessed by CD45 staining normalized to the mean vehicle (mean percentage of CD45-positive cells in the vehicle: 30.05%), with representative H&E and CD45-stained photographs. Data are mean ± SEM. n = 3, unpaired *t*-test or one-way ANOVA for multiple comparisons, **** p < 0.0001, *** p < 0.001, ** p < 0.01, * p < 0.05.

## Supplementary material

Q Exactive Hybrid Quadrupole-Orbitrap (parameters, REF), Orbitrap Fusion™ Lumos™ Tribrid™ (parameters, REF) and Orbitrap Exploris™ 480 (see below parameters).

The Vanquish Neo UHPLC System (Thermo Scientific) was coupled to an Orbitrap Exploris™ 480 mass spectrometer (Thermo Scientific), equipped with an EASY- Spray™ nanospray source (Thermo Scientific). The peptide extracts were loaded onto a 5 mm × 300 µm ID PepMap Neo Trap Cartridge (C18, 5 µm particle size, 100 Å pore size, Thermo Scientific) and separated on an analytical column (25 cm × 75 μm ID, 1.7 µm, C18 beads, Ionopticks) at a flow rate of 300 nL/min at 50°C using a multistep gradient of 3–25% mobile phase B (80% MeCN in 0.1% formic acid) for 45 minutes and 25–35% B for 15 minutes, 35–95% B for 1 minute and an 11-minute wash at 99% B. The mass spectrometer operated in positive ion mode at a 1.4 kV needle voltage, and data were acquired using Xcalibur 4.5 software in a data-dependent mode. MS scans (m/z 375–1500) were recorded at a resolution of R = 120000 (@ m/z 200), a standard AGC target, and an injection time in automatic mode, followed by a top speed duty cycle of up to 1 second for MS/MS acquisition. Precursor ions (2–6 charge states) were isolated in the quadrupole with a mass window of 2 Th and fragmented with HCD @ 30% normalized collision energy. MS/MS data were acquired with a resolution of R = 15000 (@m/z 200), a standard AGC target, and a maximum injection time in automatic mode. Selected precursors were excluded for 45 seconds.

Protein identification and label-free quantification (LFQ) were done in Proteome Discoverer 3.0. The CHIMERYS node using the prediction model inferys_2.1 fragmentations was used to identify proteins in batch mode by searching against the UniProt Homo sapiens database (82408 entries, released September 2023). Two missed enzyme cleavages were allowed for trypsin. Peptide lengths of 7–30 amino acids, a maximum of 3 modifications, charges of 2–4, and 20 ppm for fragment mass tolerance were set. Oxidation (M) and carbamidomethyl (C) were respectively searched as dynamic and static modifications by the CHIMERYS software. Peptide validation was performed using the Percolator algorithm (REF) and only “high confidence” peptides were retained corresponding to a 1% false discovery rate at the peptide level. Minora feature detector node (LFQ) was used along with the feature mapper and precursor ions quantifier. The normalization parameters were selected as follows: (1) Unique peptides, (2) Precursor abundance based on intensity, (3) Normalization mode: total peptide amount from *Homo sapiens*, (4) Protein abundance calculation: summed abundances, (5) Protein ratio calculation: pairwise ratio based and (6) Missing values were replaced with random values sampled from the lower 5% of detected values. Quantitative data were considered for master proteins and quantified by a minimum of 2 unique peptides.

